# Temporal variation of patch connectivity determines biodiversity recovery from recurrent disturbances

**DOI:** 10.1101/2022.01.02.474736

**Authors:** Claire Jacquet, François Munoz, Núria Bonada, Thibault Datry, Jani Heino, Franck Jabot

## Abstract

Understanding the capacity of ecological systems to withstand and recover from disturbances is a major challenge for ecological research in the context of environmental change. Disturbances have multi-scale effects: they can cause species extinctions locally and alter connectivity between habitat patches at the metacommunity level. Yet, our understanding of how disturbances influence landscape connectivity remains limited. To fill this gap, we develop a novel connectivity index that integrates the temporal variation of patch connectivity induced by disturbances, which can be applied to any spatially-structured habitat. We then combine this index with a metacommunity model to specifically investigate biodiversity recovery from drying events in river network metacommunities. We demonstrate that patch connectivity explains variations of species richness between groups of organisms with contrasting dispersal modes and captures the effect of drying intensity (i.e., fraction of patches that dry-up) and drying location on community recovery. As a general rule, loss of patch connectivity decreases community recovery, regardless of patch location in the river network, dispersal mode, or drying intensity. Local communities of flying organisms maintained higher patch connectivity in drying river networks compared to organisms with strictly aquatic dispersal, which explained the higher recovery capacity of this group from drying events. The general relationship between patch connectivity and community recovery we found can be applied to any spatial network subject to temporal variation of connectivity, thus providing a powerful tool for biodiversity management in dynamic landscapes.

## INTRODUCTION

Most ecological systems are exposed to a wide range of environmental fluctuations and disturbances, both natural and human induced. These disturbances vary in intensity, frequency, duration and spatial extent (Sousa 1984; Miller *et al*. 2011; Donohue *et al*. 2016; Thom & Seidl 2016). Pulse disturbances are a specific type of disturbances, which cause a sudden increase in population mortality and alter community composition for a limited period of time (Bender *et al*. 1984, Jentsch & White 2019). Such disturbances can have multiple origins, such as wildfires (Turco *et al*. 2018), forest cutting (Nordén *et al*. 2009), flooding (Woodward *et al*. 2016; Rolls *et al*. 2018) or drying events (Messager *et al*. 2021; Sarremejane *et al*. 2021). They are likely to increase in frequency and intensity due to climate change in most regions of the world (Coumou *et al*. 2012; Harris *et al*. 2018), questioning the capacity of ecological systems and organisms to withstand and recover from recurrent pulse disturbances (Jacquet *et al*. 2020).

Although some organisms have developed resistance traits to cope with disturbance events (e.g., seed banks, eggs or resting stages that survive in extreme conditions), much of the resilience of ecological communities, defined as their capacity to recover a pre-disturbance state after a disturbance, depends on species recolonization from undisturbed neighbouring habitats (Tonkin *et al*. 2018; Van Looy *et al*. 2019). Metapopulation theory has outlined the key role of species dispersal capabilities and landscape structure to determine metapopulation persistence and species extinction probability (Hanski & Ovaskainen 2000). Furthermore, works on graph theory showed that patch connectivity is positively related to local species richness, both in random networks (Bunn *et al*. 2000; Urban & Keitt 2001; Limdi *et al*. 2018), lattice landscape (Laroche *et al*. 2020), as well as dendritic networks (Carrara *et al*. 2014; Tonkin *et al*. 2014, 2018).

Dispersal ability strongly influences how organisms perceive and are affected by landscape connectivity (Taylor *et al*. 1993). This has motivated the development of so-called ecologically-scaled landscape indices that incorporate the dispersal ability of an organism into the computation of landscape connectivity (Vos *et al*. 2001). In this setting, landscape connectivity is not only a purely geographical proxy of dispersal, but rather an ecological measure acknowledging the interaction between landscape geographical features and organisms’ movement capabilities. Such ecologically-scaled landscape metrics have been shown to have better abilities to predict metacommunity patterns (Laroche *et al*. 2020) and dynamics (Lalechère *et al*. 2017; Cunillera-Montcusi *et al*. 2021), and they are considered pivotal for designing efficient landscape conservation strategies (Meurant *et al*. 2018). A step forward is to acknowledge the additional role played by disturbances in shaping landscape connectivity. Indeed, disturbances can alter connectivity between patches by affecting dispersal routes, propagule sources or propagule recruitment. Taking into account a temporal variation in connectivity should thus provide additional insights into biodiversity maintenance (Martensen *et al*. 2017; Uroy *et al*. 2021). In particular, disturbances can vary in intensities, durations, frequencies and spatial locations (Donohue *et al*. 2016; Kéfi *et al*. 2019) with potentially different effects on landscape connectivity.

Drying river networks are suitable systems to study the resilience of metacommunities to recurrent disturbances. They are ubiquitous throughout the world, as large proportion as 51-60% of global stream length is dry for at least one month each year (Messager *et al*. 2021), and their extent is likely to increase with ongoing global environmental change in most regions of the world (Harris *et al*. 2018; Spinoni *et al*. 2018). Drying events have a two-fold effect on riverine metacommunities: first, they cause sudden mortality in patches that dry-up (Soria *et al*. 2017), and, second, they temporarily increase patch isolation, thus modifying patch connectivity and metacommunity networks (Gauthier *et al*. 2021). Hence, a metacommunity approach is required to fully understand biodiversity dynamics in drying river networks (Cid *et al*. 2020).

Organisms inhabiting river networks are characterized by various dispersal modes (e.g., swimming, drifting or flying) and their perception of landscape connectivity strongly differ among them (e.g., Kärnä *et al*. 2015). The dispersal of aquatic organisms is constrained by watercourse distances between local patches and, therefore, by river dendritic structure, while the dispersal of organisms with a flying adult stage (e.g., most insects) is limited by the overland distances between local patches (Cañedo-Arguelles *et al*. 2015; Heino *et al*. 2017; Gauthier *et al*. 2021; Larsen *et al*. 2021). Although many empirical studies on riverine metacommunities outlined that each dispersal mode can be associated with specific biodiversity patterns and resilience to drying events (e.g., Sarremejane *et al*. 2017, 2020; Gauthier *et al*. 2021), we still lack a mechanistic understanding of the link between dispersal mode, patch connectivity and biodiversity recovery from drying events in dendritic river networks.

Here, we first propose a novel index of connectivity that integrates species dispersal capabilities and the temporal variation of patch connectivity induced by recurrent disturbances, which can be applied to any spatially-structured habitat. Second, we design a model of river metacommunity to generate controlled biodiversity patterns in perennial conditions, as well as biodiversity trajectories following various scenarios of drying intensity, duration and location. We use this controlled simulation setting to test the ability of our connectivity index to explain variations in species richness and community recovery from drying events (i) between local patches of a given simulation and (ii) between simulated scenarios. We specifically address the following questions: 1) Can differences in species-specific patch connectivity lead to contrasting biodiversity patterns between groups of organisms with different dispersal modes? 2) What are the effects of drying events of varying intensity, duration and location on average species richness and average patch connectivity? 3) Can local biodiversity recovery be predicted by patch connectivity?

## MATERIAL AND METHODS

### 1 Patch connectivity in spatial networks subject to disturbances

We defined patch connectivity as the probability for a local community to receive individuals from all other communities of the metacommunity (Bunn *et al*. 2000; Urban & Keitt 2001, Laroche *et al*. 2020). Here, patch connectivity depends on (i) landscape structure, (iii) species dispersal type, and (iii) intensity, frequency, duration and location of disturbances, which cause temporary alterations of patch connectivity (Fig. 1). At a given time *t*, patch connectivity *C*_*j*_ (*t*) quantifies the sum of link’s weights between patch *j* and all patches of the network:

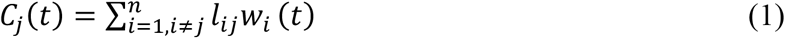

**Figure 1:**
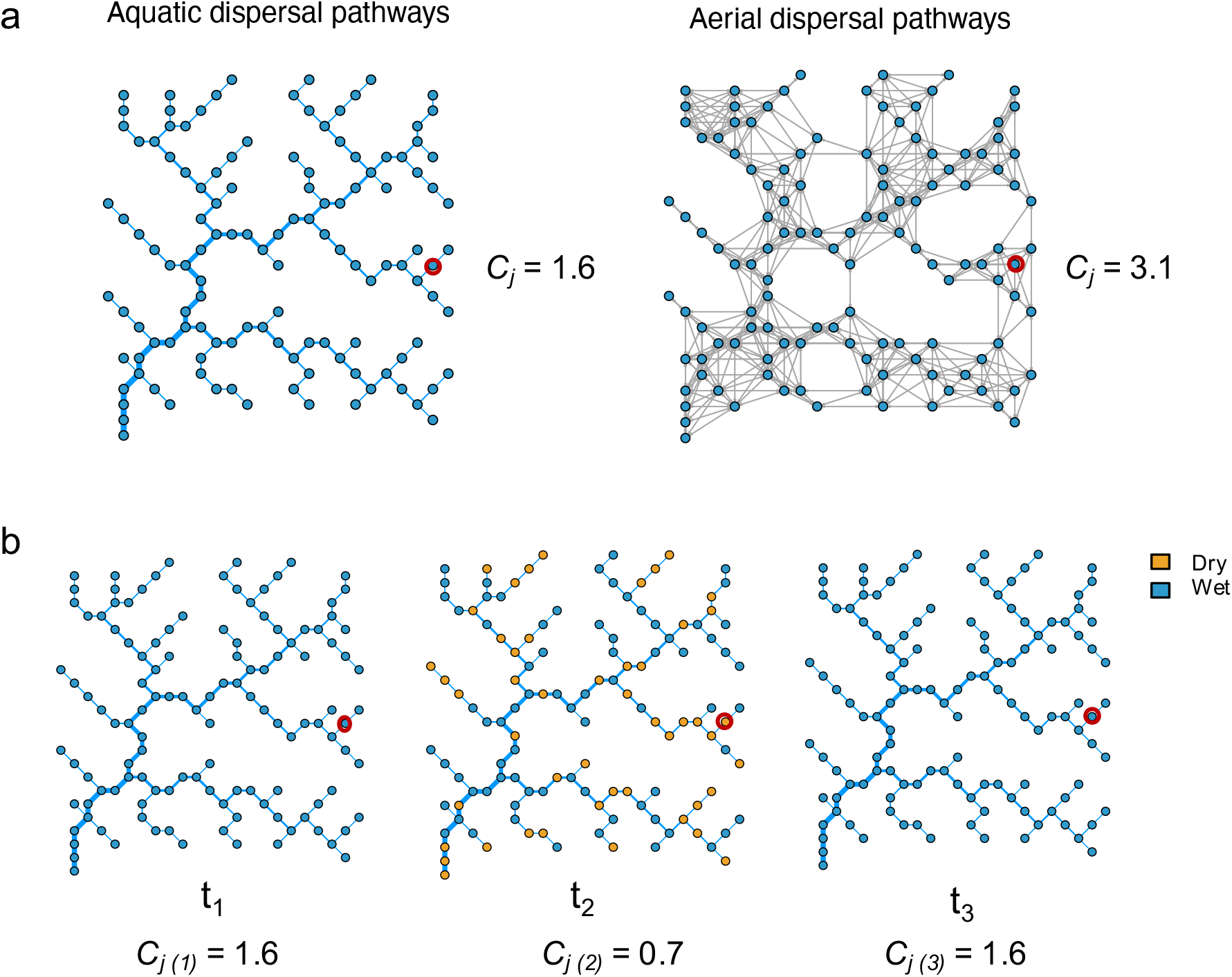
Measuring patch connectivity in a drying river network. a) Connectivity *C*_*j*_ of a local patch *j* (highlighted in red) for swimming (left) and flying (right) dispersers in a virtual river network. Line width reflects river size on the left panel, and lines connect patches located at a maximum overland distance of 2 km on the right panel. b) Illustration of the temporal variation of patch connectivity induced by drying events for a local patch (highlighted in red). Orange patches correspond to dry patches. In this example, a drying event occurs at t = 2 with an intensity of 0.4, a duration of 1 week, and a random location of dry patches.

Where *n* is the number of patches in the network, *w*_*i*_(*t*) is the viability of patch *i* at time *t* (ranging from 0 to 1, 0 if fully disturbed, 1 if undisturbed) and *l*_*ij*_ is the probability weight of the link between patches *i* and *j*, which decreases according to the formula exp(-*d*_*ij*_/*λ*), where *d*_*ij*_ is the distance between patches *i* and *j* (e.g., watercourse or overland distance for swimming or flying organisms, respectively). The scaling parameter *λ* corresponds to the average dispersal distance of the target species or dispersal group. Importantly, disturbed patches *i* at a given time *t* contribute less to *C*_*j*_ (*t*) as they contain less individuals (*w*_*i*_(*t)* < 1) while fully disturbed patches *i* at a given time *t* do not contribute at all to *C*_*j*_ (*t*) as they are empty (*w*_*i*_(*t)* = 0). Consequently, disturbance intensity, defined as the fraction of disturbed patches, necessarily decreases patch connectivity (Fig. 1b).

Connectivity *C*_*j*_ of patch *j* under a given disturbance scenario is obtained by averaging all *C*_*j*_(*t*) over time:

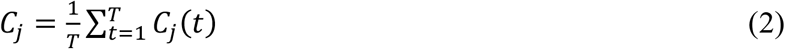

Where *T* corresponds to the time period studied (e.g., one year). Hence, disturbance duration and frequency have a negative effect on patch connectivity *C*_*j*_. We further defined patch connectivity loss as the connectivity in disturbed spatial networks relative to control (undisturbed) conditions, that is (*C*_*Control*_ - *C*_*Disturbed*_)*/(C*_*Control*_ +1), where *C*_*Disturbed*_ and *C*_*Control*_ correspond to connectivity metrics in disturbed and undisturbed spatial networks, respectively.

## 2 Metacommunity dynamics in dendritic river networks

### 2.1 Dendritic river network

We adapted the individual-based simulation algorithm of metacommunity dynamics in discrete time of Jabot *et al*. (2020) to riverine landscapes. We generated a virtual river network (so-called optimal channel network, OCN) with the R-package *OCNnet* (Carraro *et al*. 2020, 2021). OCNs are structures that reproduce the topological connectivity and scaling features of real river networks. We built a virtual river network on a square lattice spanning an area of 156.25 km^2^ (25 × 25 cells with 0.5 km cell side), with an outlet located close to the bottom left corner of the lattice. We used a threshold area of 1.25 km^2^ (5 cells) to partition the river network into *n =* 124 patches, which are illustrated by blue points in Fig. 1 (see Carraro *et al*. (2020, 2021) for detailed information on OCNs generation). We assumed that environmental conditions in perennial river networks are homogeneous across the network, and thus specifically focused on the combined effects of landscape structure and species dispersal on biodiversity dynamics. This assumption is particularly suited to model biodiversity dynamics in systems experiencing recurrent disturbance events, such as drying events or floods (Datry *et al*. 2016, 2017; Sarremejane *et al*. 2021).

### 2.2 Community dynamics

We considered a regional species pool of 100 species, each species having the same regional frequency (Table S1). Each local community had a carrying capacity of *K* = 5000 individuals, therefore local populations of species were composed of 50 individuals on average (Table S1). Both dispersal-driven regional processes and local demographic processes (births and deaths) drove biodiversity dynamics. In each local patch and at each time step, we simulated four processes taking place sequentially: (i) mortality, (ii) reproduction, (iii) dispersal, and (iv) establishment, which accounts for competition for resources (fixed carrying capacity *K* in local patches). At each time step, we simultaneously updated community composition in all patches.

#### (i) Mortality

We modelled individual mortality in the metacommunity at each time step with a Bernoulli draw with fixed mortality probability *d*. In addition, the occurrence of drying events also influenced patch viability and therefore individual mortality. We added a component to the model of Jabot *et al*. (2020) to account for pulse increase of individual mortality induced by drying events. At a given time *t*, a local patch can be dry or wet. A drying event occurring at time *t* in patch *i* causes the death of all organisms in patch *i* (i.e., all species densities are set to zero), therefore *w*_*i*_(*t)* = 0 if patch *i* is dry and *w*_*i*_(*t)* = 1 otherwise (eq. 1). Hence, we assumed that organisms do not have resistance strategies to cope with drying events (e.g., resting stages, adaptations to survive in sediments) and are not able to leave a patch when it becomes dry.

#### (ii) Reproduction

Individuals produce juveniles at a constant rate *r*, so that the number of juveniles produced by each individual during one step is a Poisson draw with parameter *r*. In the following, we considered equal mortality and reproduction rates (*r = d*) for sake of simplicity.

#### (iii) Dispersal

We considered three specific modes of dispersal that are commonly observed in riverine communities: (i) drifting, with downstream flow-directed dispersal along the waterway (e.g., plant seeds and planktonic larvae), (ii) swimming, with bidirectional dispersal along the waterway (e.g., fishes), and (iii) flying, with overland dispersal (insects with flying adult stages). For all dispersal modes, a proportion (1-*m*) of the individuals remained in the local patches, while a proportion *m* dispersed to neighbouring patches. For drifting dispersal, individuals dispersed to the patch situated downstream only (i.e., maximum drifting distance equals 0.5 km). For swimming dispersal, individuals dispersed to patches that are connected by water flow, which could be located upstream or downstream (i.e., maximum swimming distance equals 0.5 km). For flying dispersal, individuals dispersed to the patches situated closer than a threshold overland distance of 2 km (Fig. 1a), corresponding to a maximum flying distance that has been reported in the literature for mayflies (Ephemeroptera) (Kovats et al. 1996). Note that species in other groups of aquatic insects, such as dragonflies (Odonata) and caddisflies (Trichoptera), can show larger dispersal distances than mayflies.

As illustrated in Fig. 1a, the resulting dispersal pathways strongly differed among aquatic and aerial dispersal groups within the river network. Drying events also impacted individual dispersal and generated sudden decreases in patch connectivity (Fig. 1b). If a drying event occurred at time step *t* in patch *i*, no individuals dispersed to neighbouring patches and the individuals coming from neighbouring patches did not survive (i.e., all individuals are set to zero in patch *i*). We used flow directed, watercourse and overland distances to compute the *d*_*ij*_ of eq. (1) for drifting, swimming and flying organisms, respectively. We used an average dispersal distance *λ* = 0.6 km for aquatic organisms and *λ =* 1.19 km for aerial organisms, which accounts for dispersal to diagonal cells (the distance between two neighbouring cells is 0.5 km except for the ones in diagonal, where the distance is 0.71 km).

#### (iv) Establishment

Each patch had a fixed carrying capacity of *K* individuals. The number of recruited individuals *N(t)* in a patch at time *t* followed a Poisson distribution with mean equal to *K–N(t)*, where *N(t)* was the number of surviving individuals in the patch after the mortality step. No individual was recruited whenever *N(t)* was larger than *K*. The number *N*_*j*_*(t)* of recruited individuals of each species *j* followed a multinomial draw with probabilities proportional to the numbers of individuals of each species *j* reaching the focal patch, including local offspring.

### 3 Drying scenarios

At each time step, a regional drying event could occur and was defined by (i) its intensity *I*, that is the fraction of patches that dried-up, (ii) its duration *D*, that is the number of consecutive time steps with drying conditions, and (iii) its location *L* (Table S1). We tested three distinct scenarios regarding drying location: (i) random distribution of dry patches, (ii) patches located upstream being preferentially subject to drying, (iii) patches located downstream being preferentially subject to drying. (e.g., due to combined effects of drought and water extraction).

### 4 Simulations

We initialized the metacommunity with a multinomial draw of *K* = 5000 individuals from the regional pool in each patch. After a burn-in phase of 1000 steps, we modelled drying events occurring yearly over 10 years, corresponding to 520 weekly steps. We kept all biological parameters constant (*m, d, r, K*) and varied dispersal type (drifting, swimming and flying), drying intensity (0.2, 0.4, 0,6 and 0.8), drying duration (1-6 months) as well as drying location (upstream, random or downstream). This yielded a total of 219 independent simulations, including three simulations under perennial conditions (Table S1). We recorded the composition of all patches at every step of the simulations (520 steps). We measured community recovery at each time step as the fraction of species present in drying river networks relative to perennial conditions, that is SR_Drying_/SR_Perennial_, where SR_Drying_ and SR_Perennial_ correspond to species richness in drying and perennial river networks, respectively. We also calculated local species richness and community recovery at the end of the simulations, which were averaged over the last 20 steps.

## RESULTS

### Spatial distribution of species richness in perennial river networks

Local species richness was higher in downstream patches for drifting organisms (Fig. 2a), while local richness was higher in central patches for swimming organisms (Fig. 2b). Local species richness was high in all patches for flying organisms (Fig. 2c). The average local species richness of drifting organisms was markedly lower (average SR = 40), than the one of swimming (average SR = 88) or flying organisms (average SR = 98).

**Figure 2:**
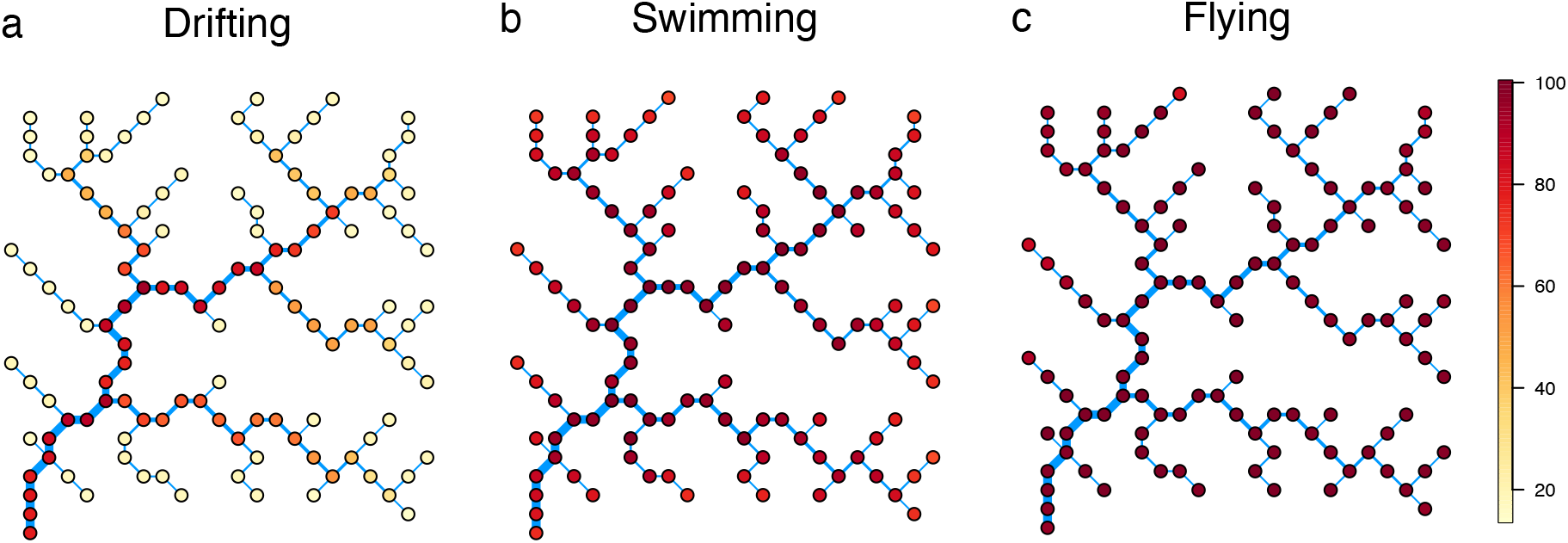
Spatial distribution of species richness (SR) in a virtual river network under perennial conditions for three dispersal modes: a) drifting (range: 14-94), b) swimming (range: 67-100) and c) flying (range: 83-100). SR values are averaged over the last 20 time-steps of the simulations. The river network is composed of 124 habitat patches and spans an area of 156.25 km^2^ (25×25 cells, cell’s side = 0.5 km). The river mouth is located at the bottom left corner of the lattices, the line width reflects river size, which increases toward the downstream direction.

### Relationship between patch connectivity and species richness

Patch connectivity allowed us to explain variations in local species richness both among different dispersal modes and among patches for a given dispersal mode (Fig. 3). Patch connectivity was systematically lower for aquatic dispersal modes compared to aerial mode: *C* ranged from 0 to 2.2 for drifting, from 0.8 to 3.1 for swimming, and from 0.9 to 5.4 for flying organisms. This ranking of patch connectivity among dispersal modes resulted in the same ranking in terms of local species richness (see the different colour groups in Fig. 3). Within each dispersal type, we further observed a positive relationship between patch connectivity and species richness. This relationship was not linear, but reached an asymptote around *C* = 3. Therefore, the correlation between patch connectivity and species richness was higher for aquatic organisms compared to aerial organisms (Pearson’s correlation, drifting: r = 0.79, swimming: r = 0.86, flying: r = 0.57, all p-values < 10^−12^).

**Figure 3:**
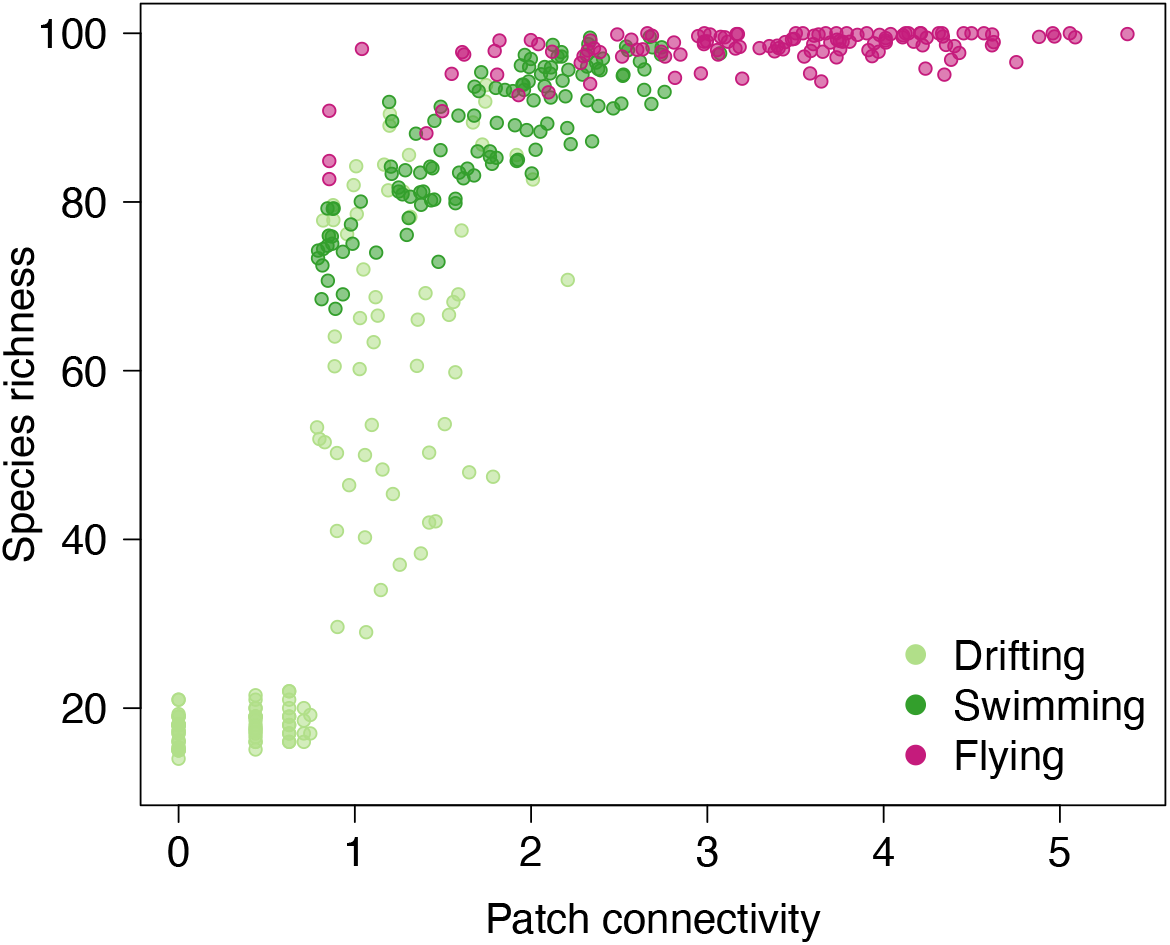
Relationship between patch connectivity and local species richness for varying dispersal modes under perennial conditions. Each point corresponds to a local patch. Points in light green, dark green, and pink correspond to local communities of drifting, swimming, and flying organisms, respectively. SR values are averaged over the last 20 time-steps of the simulations.

### Effects of drying events on average patch connectivity and species richness

Drying intensity, i.e., the fraction of patches that dry-up, had a negative effect on average patch connectivity and average species richness for all dispersal modes, regardless of the location of drying events (Fig. 4). The effect of drying intensity on average species richness was not linear, and no significant differences among drying locations were observed (Fig. 4c, Table S2). Drying duration, i.e., the number of consecutive months with dry conditions, had a negative effect on average patch connectivity (Fig. 4d). However, average species richness was not impacted by an increase in duration of drying events (Fig. 4e, Table S2).

**Figure 4:**
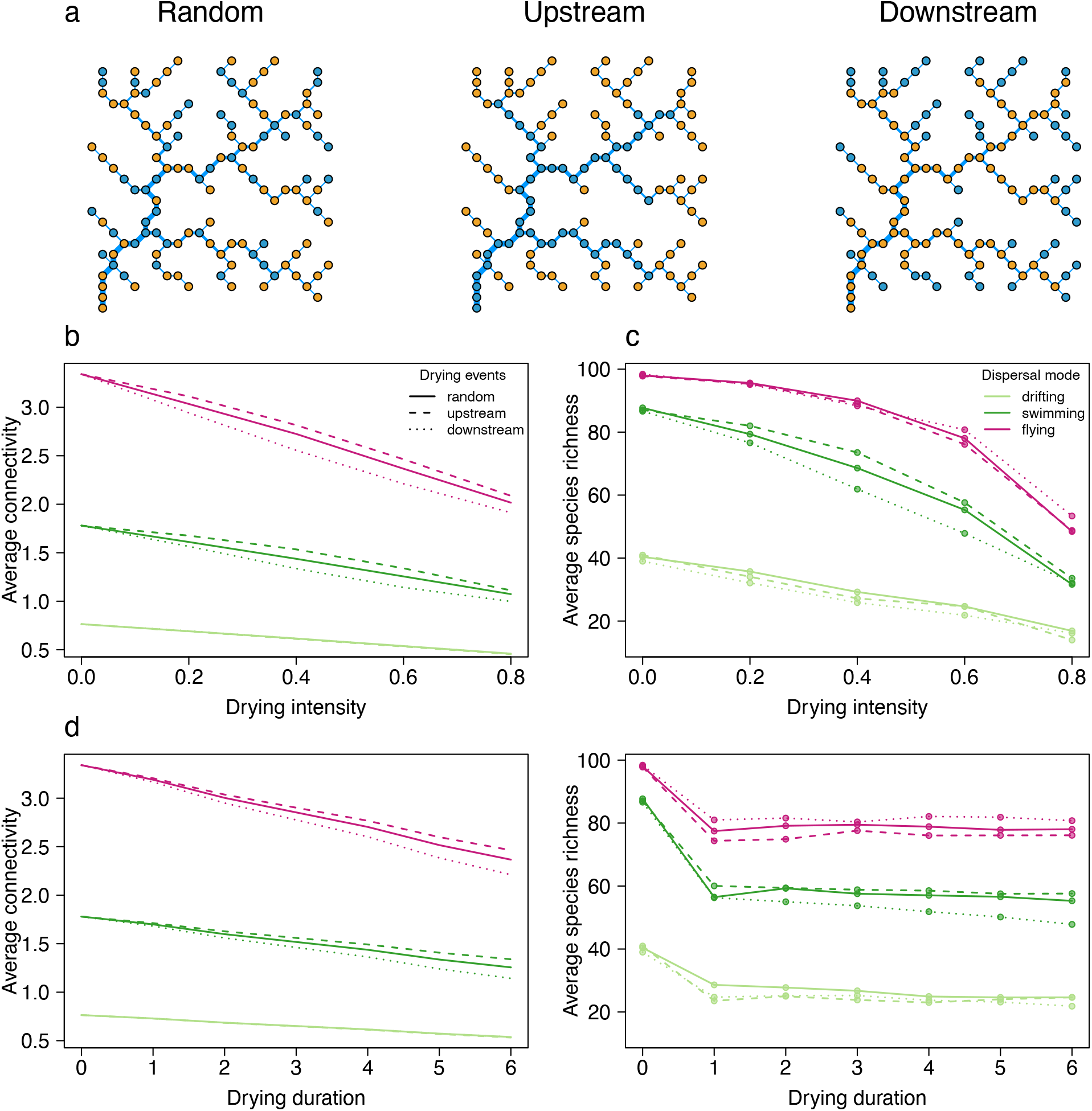
Effects of drying intensity, duration and location on average patch connectivity and average species richness for three dispersal modes. a) Location of dry patches (in orange) and perennial patches (in blue) in the river network for three scenarios of drying location: random, upstream and downstream, drying intensity *I* = 0.6 in this example. b-c) Effects of drying intensity, drying duration was fixed to 6 months. d-e) effects of drying duration, drying intensity was fixed to 0.6. Drifting, swimming and flying organisms are in light green, dark green and pink, respectively. Random, upstream and downstream drying locations are illustrated by solid, dashed and dotted lines, respectively. Results of the ANOVA testing for the effect of dispersal mode, drying intensity, location and duration on average species richness are provided in Table S2.

### Effects of drying events on community recovery

We investigated the temporal dynamics of species richness for drying intensity and duration fixed to *I =* 0.6 and *D* = 6 months. First, we observed marked variation of recovery dynamics between local communities subject to drying (Fig, 5, in orange). This variation was observed (i) between dispersal modes, (ii) between scenarios of drying location, and (iii) between local communities of the same metacommunity (i.e., within one panel of Fig. 5). Second, we observed indirect effects of drying events on the fraction of species present in perennial patches (Fig. 5, in blue). This indirect (mostly negative) effect of drying on perennial patches was also strongly heterogeneous among perennial patches.

**Figure 5:**
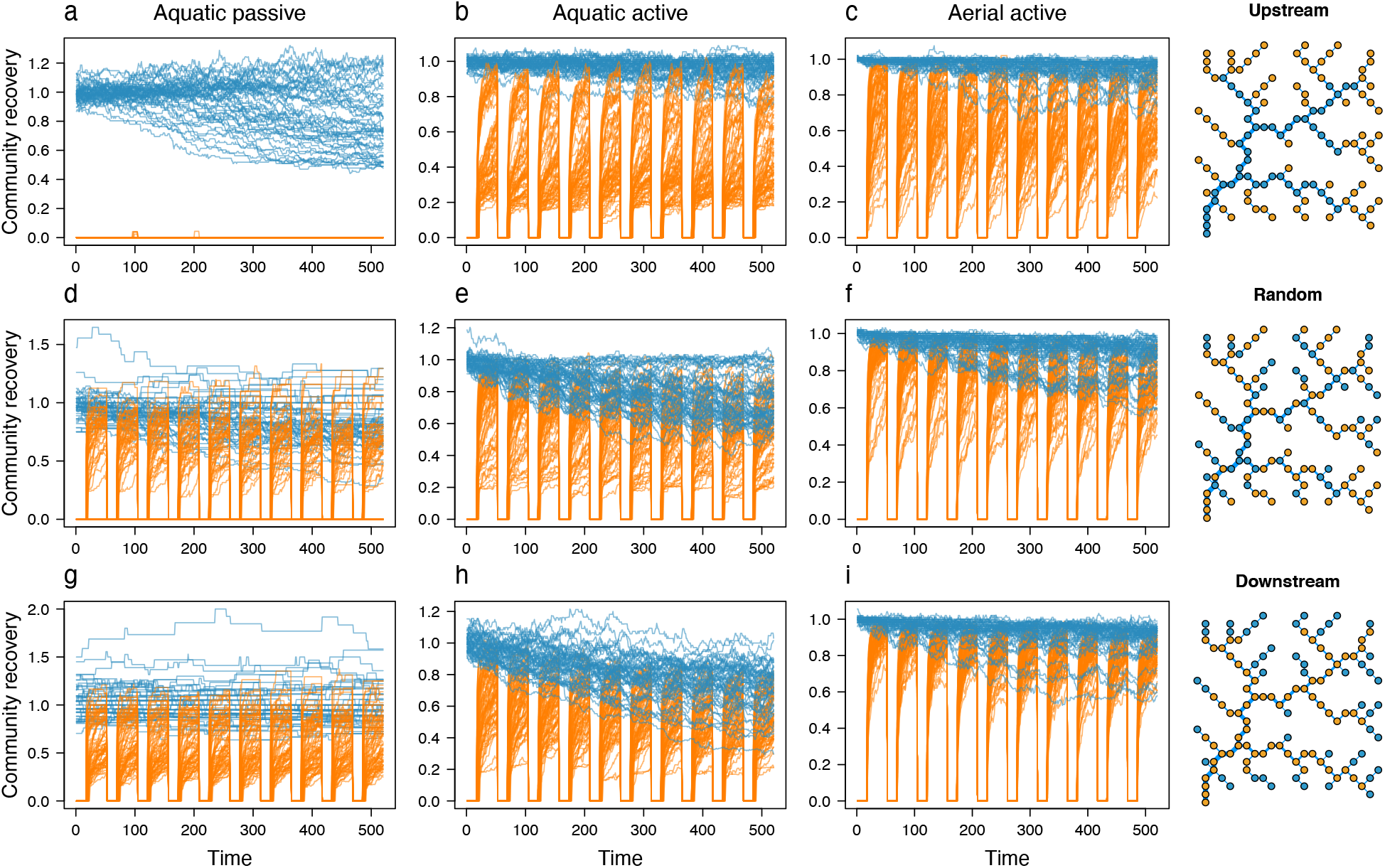
Temporal dynamics of community recovery in river networks experiencing yearly drying events for varying dispersal modes. Drying intensity: *I* = 0.6, drying duration: *D* = 6 months. Patches that dry-up are in orange and perennial patches are in blue. The river networks on the right illustrate the spatial location of dry and perennial patches (orange and blue points, respectively) for a-c) upstream, d-f) random and g-i) downstream scenarios of drying location. a, d, g) Drifting organisms, b, e, h) swimming organisms and c, f, i) flying organisms. Community recovery corresponds to local species richness relative to perennial conditions (SR_D_/SR_P_).

More precisely, drifting organisms were the most affected by drying events occurring at upstream patches (Fig. 5a), which led to the extinction of all species initially present in drying patches, and to a lower fraction of species present in several perennial patches. When drying events were located randomly or downstream, community recovery was highly variable among drying patches and some patches had even a greater richness relative to perennial conditions (Fig. 5d,g). Swimming and flying organisms were more resilient to drying events located upstream than drifting organisms (Fig. 5b,c). Swimming organisms had a lower biodiversity recovery than flying organisms when drying events were located randomly or downstream (range: 0.2-1 and 0.5-1, respectively). Flying organisms were particularly affected by drying events located upstream (Fig, 5c,f,i), while swimming organisms were similarly affected by all drying locations (Fig. 5b,e,h). We further found indirect negative effects of drying events on perennial patches (Fig. 5, in blue), which were particularly strong for swimming organisms when drying events occurred downstream or randomly (Fig. 5e,h).

### Relationship between community recovery and patch connectivity loss

Patch connectivity explained well the variations in community recovery to drying events (Fig. 6). Community final recovery was higher in patches with lower connectivity loss due to drying events, both in patches that dry up (Fig. 6a-c) and in perennial patches (Fig. 6d-f). In drying patches, community recovery of aquatic organisms decreased linearly with connectivity loss (Fig. 6a-b), while community recovery of flying organisms was little affected by connectivity loss until a threshold of 0.2 (Fig. 6c). Flying organisms had a higher proportion of local patches with high connectivity compared to other dispersal modes despite the occurrence of drying events, which explained the higher recovery ability of this group (Fig. S1). In perennial patches, community recovery did not decrease until a threshold value of connectivity loss, which differed between dispersal groups and was higher for flying organisms compared to other dispersal modes (Fig. 6d-f). Furthermore, drying events located upstream had a lower impact on the connectivity perceived by swimming and flying organisms compared to other scenarios of drying location (Fig 6e-f). The resilience of drifting organisms in perennial patches was highly variable and not well predicted by connectivity loss (Fig. 6d).

**Figure 6:**
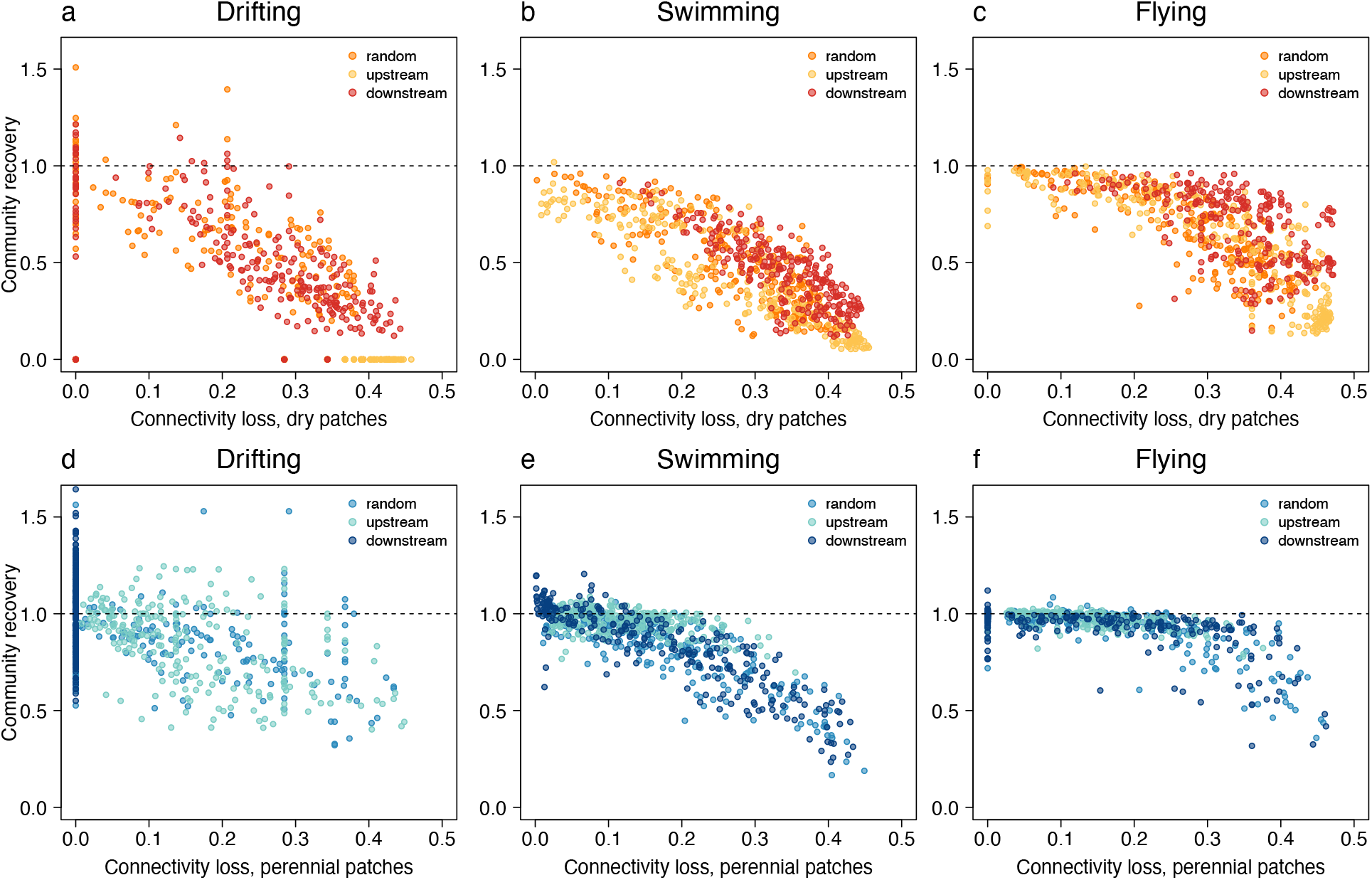
Relationship between connectivity loss and community recovery in drying and perennial patches for varying dispersal modes. Drying duration: *D* = 6 months, all drying intensities are plotted. Community recovery is averaged over the last 20 steps of the simulations. Black dotted lines indicate full recovery of species richness. a-c) Drying patches, points in yellow, orange and red correspond to upstream, random and downstream location of drying events, respectively. d-f) Perennial patches (not subject to drying events), points in turquoise, blue and dark blue correspond to upstream, random and downstream location of drying events, respectively.

## DISCUSSION

### Measuring patch connectivity in dynamic landscapes

The recognition that community composition in a habitat patch is affected by its surroundings is a central tenet of landscape ecology and has spurred the development of a myriad of connectivity indices that integrate various facets of the dispersal process, including the variability of dispersal modes and capabilities (Wilson 1992; Vos *et al*. 2001; Baranyi *et al*. 2011). However, the equal recognition that patch connectivity can be temporally variable due to disturbances is strikingly lacking. In the present work, we helped fill this gap by developing a novel connectivity index that integrates the temporal variation of patch connectivity. We demonstrated that this patch connectivity index (i) explained between-patch variations in species richness in both perennial (Fig. 3) and drying river networks (Fig. S1), (ii) also explained variations in species richness among organisms with contrasting dispersal modes (Fig. 3), and (iii) captured the effect of drying intensity and location on community recovery (Fig. 6).

Beyond its predictive power, our index of patch connectivity provides an integrated understanding of previously unrelated observations. Hence, the metacommunity model consistently reproduced the biodiversity patterns reported in theoretical and empirical studies on perennial riverine metacommunities (Fig. 2), showing that species and allelic richness of drifting organisms are highest at confluences and in downstream patches (Fagan 2002; Fernandes *et al*. 2004; Muneepeerakul *et al*. 2008; Liu *et al*. 2013), while they are highest for swimming organisms in patches occupying a central position in the river network (Economo & Keitt 2010; Carrara *et al*. 2014; Paz-Vinas & Blanchet 2015; Sarremejane et al. 2017). These apparently contrasting patterns are reconciled when framed in terms of connectivity: species richness consistently increases with connectivity for the three dispersal modes considered (Fig. 3). Similarly, variations of community recovery among local patches and simulated scenarios have a straightforward and unified explanation related to connectivity loss: connectivity loss decreases community recovery, regardless of the dispersal mode, drying intensity, or patch location (Fig. 6).

### Predicting the impact of recurrent disturbances on biodiversity recovery

Predicting how ecological communities will respond to the increased intensities and frequencies of disturbances associated with climate change is particularly challenging, since the different characteristics of disturbance events (duration, intensity, frequency, spatial location) are known to have different impacts on community recovery and species richness (Jacquet & Altermatt 2020; Jacquet *et al*. 2020). For instance, Cunillera-Montcusi *et al*. (2021) demonstrated that wildfire size and intensity have qualitatively different impacts on the recovery of macroinvertebrate communities in freshwater ponds. Similarly, we found that increasing drying intensity altered average species richness, while drying duration had almost no effect, echoing some recent empirical findings (e.g., Crabot et al. 2021). Furthermore, we showed that local species richness in undisturbed patches also decrease with connectivity loss once a threshold value of connectivity loss is reached. This result is corroborated by the empirical findings of Horváth *et al*. (2019), who used a dataset on invertebrate species in ponds spanning six decades of habitat loss to demonstrate that habitat loss also translates into species loss within the remaining habitats. Using patch connectivity as a common currency is promising to gain general insights on the ecological impact of disturbances, which are expected to be more intense and more frequent in the future (Coumou *et al*. 2012; Harris *et al*. 2018).

For the specific case of drying river networks, our study suggests some qualitative rules of thumb. First, spatial location of drying events has a comparatively second order effect on community recovery (except for drifting organisms, see Fig. 5a), which is consistent with empirical observations on macroinvertebrate metacommunities (Datry *et al*. 2014; Leigh & Datry 2017), although downstream-drying rivers show a higher resilience in some cases (Crabot *et al*. 2021). Second, drying intensity gradually alters biodiversity recovery and this effect accelerates once a threshold of connectivity loss is reached (Fig. 4c, Fig. 6), which can be expected in river networks where most of the reaches (> 80%) are intermittent. To our knowledge, there is no empirical data that tested this pattern yet, which represents a key research gap for predicting the effects of increased river network fragmentation on biodiversity. Third, organisms with various dispersal modes are likely to be differently affected by increasing drying intensity: drifting organisms being impacted first and flying organisms being impacted last (Fig. 6). This corroborates recent empirical results (e.g., Sarremejane *et al*. 2017; Gauthier *et al*. 2021; Crabot *et al*. 2021) and calls for integrating dispersal abilities of organisms when assessing the effects of river network fragmentation by drying.

### Model assumptions and perspectives

Our metacommunity model assumes that all environmental variables are similar between local patches. This is, however, an oversimplification as river networks are known to exhibit a wide range of variation of patch size and environmental conditions along the longitudinal gradient (Vannote *et al*. 1980; Carrara *et al*. 2014) as well as between different headwater sites (Heino *et al*. 2013; 2015), which are both exacerbated in drying river networks (Datry *et al*. 2016; Sarremejane *et al*. 2021). Here, our aim was to focus on spatial dynamics, which is central in landscapes subject to disturbances (Datry *et al*. 2014; Gauthier *et al*. 2021; Sarremejane *et al*. 2021). Assessing the additional contribution of environmental heterogeneity on metacommunity dynamics in such periodically disturbed systems should provide complementary insights (Holyoak *et al*. 2020, Jabot *et al*. 2020).

The model further assumes that organisms do not have resistance strategies to cope with drying events such as resting stages or adaptations to survive in the sediments or in the hyporheic zone (Datry *et al*. 2014). In the model predictions, drying events with a duration ranging from 1 to 6 months had similar effects on average species richness, suggesting that drying events of short duration are as detrimental to organisms as longer ones. Interestingly, taking into account resistance processes may trigger an effect of drying duration on metacommunity dynamics. Since resistance to drying conditions is likely to decrease with drying duration, we should observe a negative effect of drying duration on community recovery (Datry 2012; Datry *et al*. 2014; Crabot *et al*. 2021). Such a prediction is consistent with the empirical observation of a rapid recovery (within weeks) of local species richness after a short 1-week drought (Vander Vorste *et al*. 2016; Sarremejane *et al*. 2021) and a negative effect of drying duration on species richness in drying river networks (Datry 2012, Datry *et al*. 2014, Vander Vorste et al. 2016, Crabot *et al*. 2021).

### Significance of the results beyond drying river networks

Many other ecological systems are subject to recurrent disturbances such as wildfires (Turco *et al*. 2018), forest cutting (Nordén *et al*. 2009) or flooding (Woodward *et al*. 2016). Human-managed habitats also face recurrent disturbances linked to agricultural activities such as tillage, fertilisation, crop harvest and rotation (Gaba *et al*. 2018), and silvicultural activities such as forest thinning or cutting (Senf & Seidl 2021). Understanding biodiversity dynamics and resilience in such recurrently disturbed systems requires developing quantitative frameworks in which transient dynamics following disturbance events is the rule rather than the exception (Pickett & White 1985; Fukami & Nakajima 2011). Our contribution provides an operational connectivity measure that encapsulates the influence of disturbances on temporally variable landscape connectivity. We demonstrated its utility for understanding biodiversity dynamics in drying river networks, but this tool can also be used in a number of other ecological systems to analyse the contribution of patch connectivity to community recovery.

## REFERENCES

Baranyi, G., Saura, S., Podani, J. & Jordán, F. (2011). Contribution of habitat patches to network connectivity : Redundancy and uniqueness of topological indices. Ecol. Indic., 11, 1301–1310.

Bender EA, Case TJ, Gilpin ME. (1984) Perturbation experiments in community ecology: theory and practice. Ecology 65, 1–13.

Bunn, A.G., Urban, D.L. & Keitt, T.H. (2000). Landscape connectivity : A conservation application of graph theory. J. Environ. Manage., 59, 265–278.

Cañedo-Argüelles, M., Boersma, K.S., Bogan, M.T., Olden, J., Phillipsen, I., Schriever, T.A., et al. (2015). Dispersal strength determines meta-community structure in a dendritic riverine network. J. Biogeogr., 42, 778–790.

Carrara, F., Rinaldo, A., Giometto, A. & Altermatt, F. (2014). Complex Interaction of Dendritic Connectivity and Hierarchical Patch Size on Biodiversity in River-Like Landscapes. Am. Nat., 183, 13–25.

Carraro, L., Bertuzzo, E., Fronhofer, E.A., Furrer, R., Gounand, I., Rinaldo, A., et al. (2020). Generation and application of river network analogues for use in ecology and evolution. Ecol. Evol., ece3.6479.

Carraro, L., Bertuzzo, E., Fronhofer, E.A., Furrer, R., Gounand, I., Rinaldo, A., et al. (2021). Package “OCNet” https://CRAN.R-project.org/package=OCNet.

Cid, N., Bonada, N., Heino, J., Cañedo-Argüelles, M., Crabot, J., Sarremejane, R., et al. (2020). A Metacommunity Approach to Improve Biological Assessments in Highly Dynamic Freshwater Ecosystems. Bioscience, 70, 427–438.

Coumou, D. & Rahmstorf, S. (2012). A decade of weather extremes. Nat. Clim. Chang., 2, 491–496.

Crabot, J., Mondy, C.P., Usseglio-polatera, P., Fritz, K.M., Wood, P.J., Greenwood, M.J., et al. (2021). A global perspective on the functional responses of stream communities to flow intermittence. Ecography (Cop.)., 1–13.

Cunillera-Montcusí, D., Borthagaray, A.I., Boix, D., Gascón, S., Sala, J., Tornero, I., et al. (2021). Metacommunity resilience against simulated gradients of wildfire: disturbance intensity and species dispersal ability determine landscape recovery capacity. Ecography (Cop.)., 44, 1022–1034.

Datry, Thibault. (2012). Benthic and hyporheic invertebrate assemblages along a flow intermittence gradient: effects of duration of dry events. Freshwater Biology. 57. 563–574.

Datry, T., Larned, S. T., Fritz, K. M., Bogan, M. T., Wood, P. J., Meyer, E. I., & Santos, A. N. (2014). Broad-scale patterns of invertebrate richness and community composition in temporary rivers: Effects of flow intermittence. Ecography, 37(1), 94–104.

Datry, T., Pella, H., Leigh, C., Bonada, N. and Hugueny, B. (2016), A landscape approach to advance intermittent river ecology. Freshw Biol, 61: 1200–1213. https://doi.org/10.1111/fwb.12645

Datry, T., Corti, R., Heino, J., Hugueny, B., Rolls, R.J. & Ruhí, A. (2017). Habitat Fragmentation and Metapopulation, Metacommunity, and Metaecosystem Dynamics in Intermittent Rivers and Ephemeral Streams. In: Intermittent Rivers and Ephemeral Streams. Elsevier, pp. 377–403.

Donohue, I., Hillebrand, H., Montoya, J.M., Petchey, O.L., Pimm, S.L., Fowler, M.S., et al. (2016). Navigating the complexity of ecological stability. Ecol. Lett., 19, 1172–1185.

Economo, E.P. & Keitt, T.H. (2010). Network isolation and local diversity in neutral metacommunities. Oikos, 1–9.

Fagan, W.F. (2002). Connectivity, fragmentation, and extinction risk in dendritic metapopulations. Ecology, 83, 3243–3249.

Fernandes, C.C., Podos, J. & Lundberg, J.G. (2004). Amazonian ecology: Tributaries enhance the diversity of electric fishes. Science (80-.)., 305, 1960–1962.

Fukami, T. & Nakajima, M. (2011). Community assembly: Alternative stable states or alternative transient states? Ecol. Lett., 14, 973–984.

Gaba S. et al. (2018) Ecology for Sustainable and Multifunctional Agriculture. In:Gaba S., Smith B., Lichtfouse E. (eds) Sustainable Agriculture Reviews 28. Sustainable Agriculture Reviews, vol 28. Springer, Cham.

Smith B., Lichtfouse E. (eds) Sustainable Agriculture Reviews 28. Sustainable Agriculture Reviews, vol 28. Springer, Cham.

Gauthier, M., Le Goff, G., Launay, B., Douady, C.J. & Datry, T. (2021). Dispersal limitation by structures is more important than intermittent drying effects for metacommunity dynamics in a highly fragmented river network. Freshw. Sci., 40, 302–315.

Hanski, I. & Ovaskainen, O. (2000). The metapopulation capacity of a fragmented landscape. Nature, 404, 755–758.

Harris, R.M.B., Beaumont, L.J., Vance, T.R., Tozer, C.R., Remenyi, T.A., Perkins-Kirkpatrick, S.E., et al. (2018). Biological responses to the press and pulse of climate trends and extreme events. Nat. Clim. Chang., 8, 579–587.

Heino, J., Grönroos, M., Ilmonen, J., Karhu, T., Niva, M. & Paasivirta, L. (2013) Environmental heterogeneity and beta diversity of stream macroinvertebrate communities at intermediate spatial scales. Freshwater Science 32: 142–154.

Heino, J., Melo, A.S., Siqueira, T., Soininen, J., Valanko, S. & Bini, L.M. (2015). Metacommunity organisation, spatial extent and dispersal in aquatic systems: patterns, processes and prospects. Freshw. Biol., 60, 845–869.

Heino, J., Alahuhta, J., Ala-Hulkko, T., Antikainen, H., Bini, L. M., Bonada, N., … & Soininen, J. (2017). Integrating dispersal proxies in ecological and environmental research in the freshwater

Holyoak, M., Caspi, T. & Redosh, L.W. (2020). Integrating Disturbance, Seasonality, Multi-Year Temporal Dynamics, and Dormancy Into the Dynamics and Conservation of Metacommunities. Front. Ecol. Evol., 8, 1–17.

Horváth, Z., Ptacnik, R., Vad, C.F. and Chase, J.M. (2019), Habitat loss over six decades accelerates regional and local biodiversity loss via changing landscape connectance. Ecol Lett, 22: 1019 1027.

Jabot, F., Laroche, F., Massol, F., Arthaud, F., Crabot, J., Dubart, M., et al. (2020). Assessing metacommunity processes through signatures in spatiotemporal turnover of community composition. Ecol. Lett., 23, 1330–1339.

Jacquet, C. & Altermatt, F. (2020). The ghost of disturbance past: long-term effects of pulse disturbances on community biomass and composition. Proc. R. Soc. B. 287: 20200678.

Jacquet, C., Gounand, I. & Altermatt, F. (2020). How pulse disturbances shape size-abundance pyramids. Ecol. Lett., 23, 1014–1023.

Jentsch, A. & White, P. (2019). A theory of pulse dynamics and disturbance in ecology. Ecology, 100, e02734.

Kärnä, O.-M., Grönroos, M., Antikainen, H., Hjort, J., Ilmonen, J., Paasivirta, L. & Heino, J. (2015) Inferring the effects of potential dispersal routes on the metacommunity structure of stream insects: as the crow flies, as the fish swims or as the fox runs? Journal of Animal Ecology 84: 1342–1353.

Kéfi, S., Domínguez-García, V., Donohue, I., Fontaine, C., Thébault, E. & Dakos, V. (2019). Advancing our understanding of ecological stability. Ecol. Lett., 22, 1349–1356.

Kovats, Z.E., Ciborowski, J.J.H. & Corkum, L.D. (1996). Inland dispersal of adult aquatic insects. Fre, 36, 265–276.

Lalechère, E., Jabot, F., Archaux, F. & Deffuant, G. (2017). Non-equilibrium plant metapopulation dynamics challenge the concept of ancient/recent forest species. Ecol. Modell., 366, 48–57.

Laroche, F., Balbi, M., Grébert, T., Jabot, F. and Archaux, F. (2020) Three points of consideration before testing the effect of patch connectivity on local species richness: patch delineation, scaling and variability of metrics. bioRxiv, 640995, ver. 5 peer-reviewed and recommended by PCI Ecology. doi: 10.1101/640995

Larsen, S., Comte, L., Filipa Filipe, A., Fortin, M., Jacquet, C., Ryser, R., et al. (2021). The geography of metapopulation synchrony in dendritic river networks. Ecol. Lett., 24, 791–801.

Leigh, C. and Datry, T. 2017. Drying as a primary hydrological determinant of biodiversity in river systems: a broad-scale analysis. –Ecography 40: 487–499.

Limdi, A., Pérez-Escudero, A., Li, A. & Gore, J. (2018). Asymmetric migration decreases stability but increases resilience in a heterogeneous metapopulation. Nat. Commun., 9, 2969.

Liu, J., Soininen, J., Han, B.P. & Declerck, S.A.J. (2013). Effects of connectivity, dispersal directionality and functional traits on the metacommunity structure of river benthic diatoms. J. Biogeogr., 40, 2238–2248.

Van Looy, K., Tonkin, J.D., Floury, M., Leigh, C., Soininen, J., Larsen, S., et al. (2019). The three Rs of river ecosystem resilience: Resources, recruitment, and refugia. River Res. Appl., 35, 107–120.

Martensen, A.C., Saura, S. & Fortin, M. (2017). Spatio-temporal connectivity : assessing the amount of reachable habitat in dynamic landscapes. Methods, 8, 1253–1264.

Meurant, M., Gonzalez, A., Doxa, A. & Albert, C.H. (2018). Selecting surrogate species for connectivity conservation. Biol. Conserv., 227, 326–334.

Miller, A.D., Roxburgh, S.H. & Shea, K. (2011). How frequency and intensity shape diversity-disturbance relationships. Proc. Natl. Acad. Sci. U. S. A., 108, 5643–5648.

Muneepeerakul, R., Bertuzzo, E., Lynch, H.J., Fagan, W.F., Rinaldo, A. & Rodriguez-Iturbe, I. (2008). Neutral metacommunity models predict fish diversity patterns in Mississippi–Missouri basin. Nature, 453, 220–222.

Nordén, B., Rørstad, P.K., Magnér, J. & Götmark, F. (2019). The economy of selective cutting in recent mixed stands during restoration of temperate deciduous forest. Scand. J. For. Res., 34, 709–717.

Paz-Vinas, I. & Blanchet, S. (2015). Dendritic connectivity shapes spatial patterns of genetic diversity: a simulation-based study. J. Evol. Biol., 28, 986–994.

Pickett, S.T.A. & White, P.S. (1985). The Ecology of Natural Disturbance and Patch Dynamics. Academic Press, New York.

Rolls, R.J., Heino, J., Ryder, D.S., Chessman, B.C., Growns, I.O., Thompson, R.M. and Gido, K.B. (2018), Scaling biodiversity responses to hydrological regimes. Biol Rev, 93: 971–995. https://doi.org/10.1111/brv.12381

Sarremejane, R., Mykrä, H., Bonada, N., Aroviita, J. & Muotka, T. (2017). Habitat connectivity and dispersal ability drive the assembly mechanisms of macroinvertebrate communities in river networks. Freshw. Biol., 62, 1073–1082.

Sarremejane, R., England, J., Sefton, C.E.M., Parry, S., Eastman, M. & Stubbington, R. (2020). Local and regional drivers influence how aquatic community diversity, resistance and resilience vary in response to drying. Oikos, 129, 1877–1890.

Sarremejane, R., Stubbington, R., England, J., Sefton, C.E.M., Eastman, M., Parry, S., et al. (2021). Drought effects on invertebrate metapopulation dynamics and quasi-extinction risk in an intermittent river network. Glob. Chang. Biol., 0–2.

Sebald, J., Senf, C. & Seidl, R. (2021). Human or natural ? Landscape context improves the attribution of forest disturbances mapped from Landsat in Central Europe. Remote Sens. Environ., 262, 112502.

Soria, M., Leigh, C., Datry, T., Bini, L.M. & Bonada, N. (2017). Biodiversity in perennial and intermittent rivers: a meta-analysis. Oikos, 126, 1078–1089.

Sousa, W.P. (1984). The Role of Disturbance in Natural Communities. Annu. Rev. Ecol. Syst., 15, 353–391.

Spinoni, J., Vogt, J. V., Naumann, G., Barbosa, P. & Dosio, A. (2018). Will drought events become more frequent and severe in Europe? Int. J. Climatol., 38, 1718–1736.

Taylor, P.D., Fahrig, L., Henein, K. & Merriam, G. (1993). Connectivity Is a Vital Element of Landscape Structure. Oikos, 68, 571–573.

Thom, D. & Seidl, R. (2016). Natural disturbance impacts on ecosystem services and biodiversity in temperate and boreal forests. Biol. Rev. Camb. Philos. Soc., 91, 760–781.

Tonkin, J.D., Altermatt, F., Finn, D.S., Heino, J., Olden, J., Pauls, S.U., et al. (2018). The role of dispersal in river network metacommunities: Patterns, processes, and pathways. Freshw. Biol., 63, 141–163.

Turco, M., Rosa-cánovas, J.J., Bedia, J., Llasat, M.C., Provenzale, A., Jerez, S., et al. (2018). Exacerbated fi res in Mediterranean Europe due to stationary climate-fire models. Nat. Commun., 1–9.

Urban, D. & Keitt, T. (2001). Landscape Connectivity: A Graph-Theoretic Perspective. Ecology, 82, 1205–1218.

Uroy, L., Alignier, A., Mony, C. et al. How to assess the temporal dynamics of landscape connectivity in ever-changing landscapes: a literature review. Landscape Ecol. 36, 2487–2504 (2021).

Vannote, R.L., Minshall, G.W., Cummins, K.W., Sedell, J.R. & Cushing, C.E. (1980). The River Continuum Concept. Can. J. Fish. Aquat. Sci., 37, 130–137.

Vander Vorste, R., Malard, F. & Datry, T. (2016). Is drift the primary process promoting the resilience of river invertebrate communities? A manipulative field experiment in an intermittent alluvial river. Freshw. Biol., 61, 1276–1292.

Vos, C.C., Verboom, J., Opdam, P.F.M. & Braak, C.J.F. Ter. (2001). Toward Ecologically Scaled Landscape Indices. Am. Nat., 183.

Wilson, D.S. (1992). Complex interactions in metacommunities, with implications for biodiversity and higher levels of selection. Ecology, 73, 1984–2000.

Woodward, G., Bonada, N., Brown, L.E., Death, R.G., Durance, I., Gray, C., et al. (2016). The effects of climatic fluctuations and extreme events on running water ecosystems. Philos. Trans. R. Soc., 371, 20150274.

