## Supporting Information for "Temporal variation of patch connectivity determines biodiversity recovery from recurrent disturbances"

### SUPPLEMENTARY TABLE

**Table S1:** Model parameters used in the simulations.

| Symbol | Definition | Values |
|---|---|---|
| $n$ | Number of patches | 124 |
| $S$ | Number of species in the regional pool | 100 |
| $K$ | Carrying capacity in local patches | 5000 |
| $d, r$ | Mortality/Reproduction rate | 0.1 ($week^{-1}$) |
| $m$ | Proportion of individuals that disperse | 0.25 |
| $w$ | Dispersal mode | drifting (0), swimming (1), flying (2) |
| $d_{max}$ | Maximum dispersal distance | aquatic: 0.5 (km), aerial: 2 (km) |
| $R_f$ | Regional frequency | 1/$S$ for all species |
| $I$ | Intensity of drying events | [0, 0.2, 0.4, 0.6, 0.8] |
| $D$ | Duration of drying events | [0, 1, 2, 3, 4, 5, 6] (months) |
| $L$ | Location of drying events | random, upstream, downstream |

**Table S2:** Results of the analysis of variance (ANOVA) testing for the effect of (i) dispersal mode, (ii) intensity, (iii) location and (iv) duration of drying events on average species richness (Figure 4).

| Explanatory variable | d.f. | Sum of squares | Mean of squares | F-value | p-value |
|---|---|---|---|---|---|
| Dispersal mode | 1 | 105593 | 105593 | 1731 | $< 10^{-6}$ |
| Drying intensity | 2 | 39971 | 39971 | 655 | $< 10^{-6}$ |
| Drying location | 1 | 54 | 27 | 0.4 | 0.64 |
| Duration | 1 | 0 | 0 | 0.0006 | 0.98 |
| Residuals | 219 | 13353 | 61 | | |

### SUPPLEMENTARY FIGURES

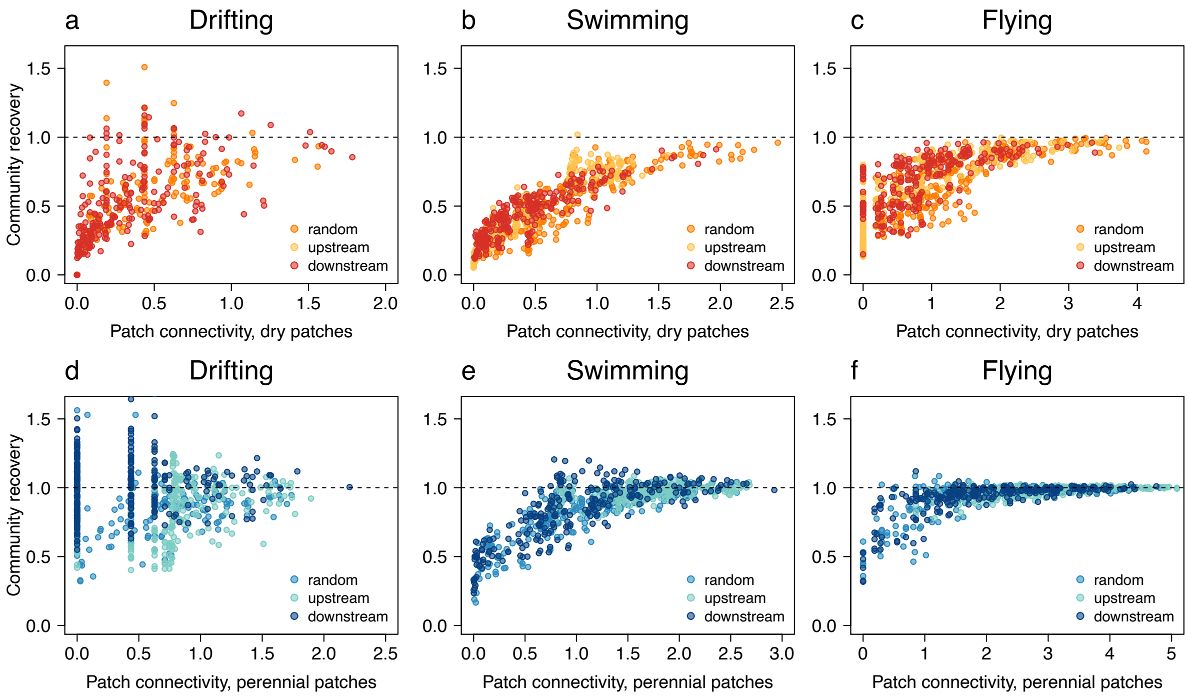


**Figure S1:** Relationship between patch connectivity (*C*) and community recovery in drying and perennial patches for varying dispersal modes. Community recovery corresponds to local species richness relative to perennial conditions averaged over the last 20 steps of the simulations. Black dotted lines indicate full recovery of species richness. a-c) Drying patches, points in yellow, orange and red correspond to upstream, random and downstream location of drying events, respectively. d-f) Perennial patches (not subject to drying events), points in turquoise, blue and dark blue correspond to upstream, random and downstream location of drying events, respectively.
